# Vaccination against antibiotic resistant gonorrhoea for men who have sex with men in England: a modelling study

**DOI:** 10.1101/692954

**Authors:** Lilith K Whittles, Peter J. White, Xavier Didelot

**Affiliations:** Department of Infectious Disease Epidemiology, School of Public Health, Imperial College London, London, UK; MRC Centre for Global Infectious Disease Analysis, School of Public Health, Imperial College London, London, UK; NIHR Health Protection Research Unit in Modelling Methodology, School of Public Health, Imperial College London, London, UK; Modelling and Economics Unit, National Infection Service, Public Health England, London, UK; School of Life Sciences University of Warwick, Coventry, UK; Department of Statistics, University of Warwick, Coventry, UK

## Abstract

**Background:** Gonorrhoea incidence is increasing rapidly: diagnoses in men who have sex with men (MSM) in England increased eight-fold 2008-2017. Concurrently, antibiotic resistance is making treatment more difficult, leading to renewed interest in a gonococcal vaccine. The MeNZB meningococcal B vaccine is partially protective, and several other candidates are in development. We modelled realistic vaccination strategies under various scenarios of antibiotic resistance and vaccine protection level and duration, to assess the impact of vaccination and examine the feasibility of the WHO’s target of reducing gonorrhoea incidence by 90% between 2016 and 2030.

**Methods:** We fitted a stochastic transmission-dynamic model, incorporating asymptomatic and symptomatic infection and heterogeneous sexual behaviour, to gonorrhoea incidence in MSM in England, 2008-17, using particle Markov Chain Monte Carlo methods. Bayesian forecasting was used to examine future scenarios, including emergence of extensively antibiotic-resistant (ABR) gonorrhoea.

**Findings:** Even in the worst-case scenario of untreatable infection emerging, the WHO target could be met by vaccinating all MSM attending sexual health clinics with a 53%-protective vaccine lasting for >6 years, or a 70%-protective vaccine lasting >3 years. A vaccine like MeNZB, conferring 30% protection for 2-4 years, could reduce incidence in 2030 by 45% in the worst-case scenario, and by 75% if >70% of ABR gonorrhoea is treatable.

**Interpretation:** Our statistically-rigorous assessment shows that even a partially-protective vaccine, delivered through a practical targeting strategy, could have substantial benefit in reducing gonorrhoea incidence in the context of an epidemic with rising antibiotic resistance.

## INTRODUCTION

The World Health Organization (WHO) classifies *Neisseria gonorrhoeae* as a priority bacterial pathogen, due to the evolution and global spread of resistance to every antibiotic historically used against it.^1^ In England there is a growing epidemic of gonorrhoea, while treatment is becoming harder, particularly among men who have sex with men (MSM).^2^ In 2019, rising azithromycin resistance in the UK has led to the drug’s removal from the previously recommended dual therapy.^3^ The remaining component, ceftriaxone, is now both the first and last line of treatment, with worrying signs that it may also fail soon.^4,5^

The threat of antibiotic resistant gonorrhoea has renewed interest in vaccine development, which is hampered by genetic variability in the gonococcus, the absence of a measurable correlate of immunity to gonorrhoea, and the lack of a suitable animal model to test vaccine candidates *in vivo*.^6^ Despite repeated efforts to develop a vaccine throughout the 20^th^ century, none of the four candidates that progressed to clinical trials were found to be effective.^7^ However, hope was recently revived when surveillance reports from Cuba, New Zealand and Norway showed a decline in gonorrhoea incidence in the years immediately following vaccination initiatives against the closely related bacterial species *Neisseria meningitidis*.^8^ A retrospective case-control study based on a million young adults in New Zealand who had received an outer-membrane vesicle meningococcal B vaccine (MeNZB) found cross-protection against *N. gonorrhoeae*, with estimated effectiveness of 31% (95% CI 21%-39%),^9^ providing the first indication that vaccination against gonorrhoea might be feasible.^7^ Additionally, there are multiple promising vaccine candidates in preclinical development stages.^8,10^

The rising antimicrobial resistance context raises the question of how best to deploy vaccines to decrease the overall burden of disease.^11–13^ In 2016, the WHO announced a global health sector strategy on sexually transmitted infections, with a target of 90% reduction in gonorrhoea incidence by 2030.^14^ For MSM in England this would correspond to <1,800 cases in 2030, based on national surveillance figures from Public Health England.^15^ Using a stochastic model of gonorrhoea epidemiology calibrated to recent incidence data, with varying future levels of antibiotic resistance, we investigated how protective and long-lasting a vaccine would need to be to reduce total incidence below the WHO target. We compared the impact and efficiency of three different vaccination strategies, studied the interplay between vaccination and antibiotic resistance levels, and quantified the effect of differing levels of vaccine uptake on the results. Finally, we assessed the potential impact of vaccines with a partially protective profile similar to MeNZB.

## METHODS

### Model structure

We developed a stochastic compartmental model to project the future course of the gonorrhoea epidemic among MSM in England under different vaccination strategies. We extended a previous model^16^ to incorporate heterogeneity in sexual risk behaviour by dividing the population into low- and high-risk groups with characteristic rates of sexual partner change. The model simulates gonorrhoea in MSM in England from 2008 to 2030. The initial size and rates of entry and exit from the sexually active population were based on projections from the Office of National Statistics.^17^ The proportion of the population in each sexual activity group and their respective annual number of partners were calibrated according to data from the third National Survey of Sexual Attitudes and Lifestyles (Natsal-3).^18^ The rate of transmission is dependent on risk group.

Following infection with gonorrhoea, individuals initially pass through a short incubation period, after which they may develop symptoms or remain asymptomatically infected (Figure 1). Gonococcal infection can occur in the rectum, pharynx and/or urethra, but surveillance data are not stratified by infection site, thus the parameters are averages for all sites. Infected individuals are treated after seeking care due to symptoms or after testing positive in sexual health screening. Treated individuals recover from the infection and become uninfected again, with the exception of a proportion of those infected with the antibiotic resistant (ABR) strain for whom treatment may fail, leading to a persistent asymptomatic infection.^16^ Recovery from asymptomatic infection eventually occurs naturally, or following antibiotic intake for unrelated conditions. Infection is assumed not to confer any natural immunity.^19,20^ The ABR strain is assumed to emerge globally in 2020 and subsequently to be imported repetitively from overseas into the highly sexually active group in England.^5^

**Figure 1.**
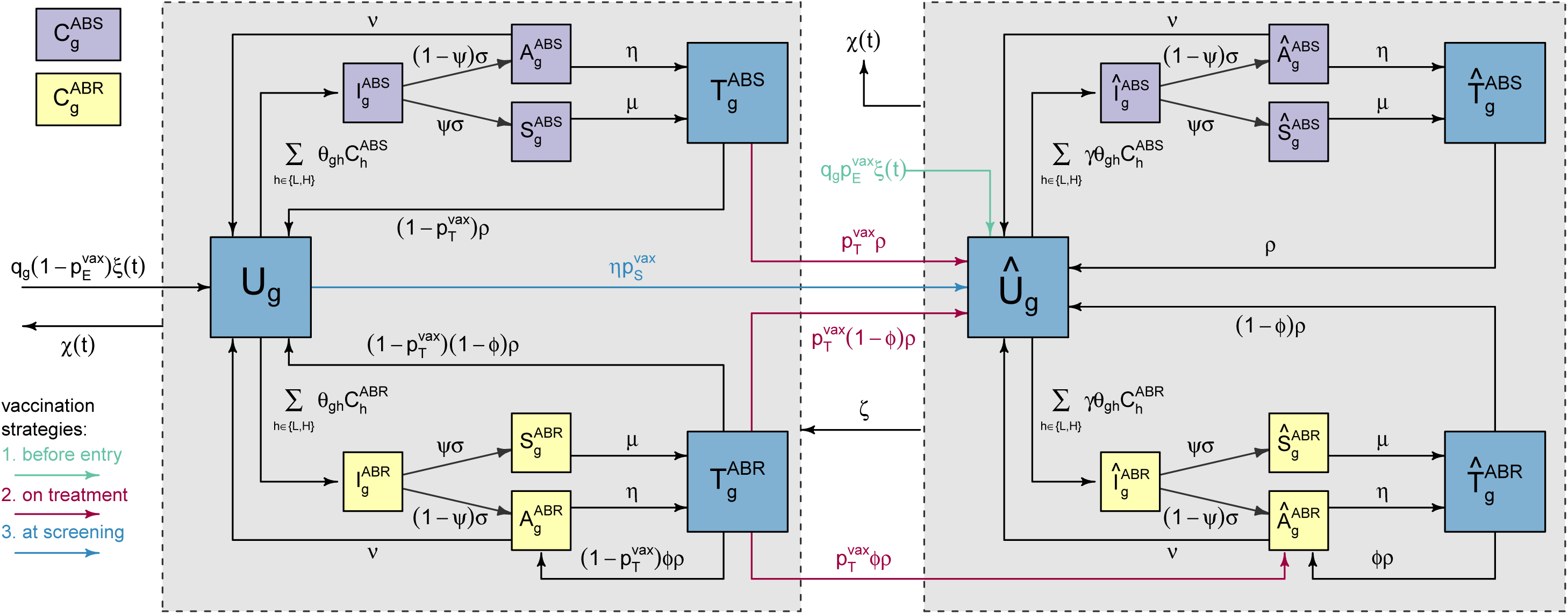
Model structure flow diagram. Uninfected individuals (*U*) in sexual activity group *g* (low or high) become infected with strain *s* (antibiotic-sensitive (ABS) or antibiotic-resistant (ABR)). Infected individuals pass through an incubating state (*I)*, before either developing symptoms (*S*) or remaining asymptomatic (*A*). Symptomatic individuals seek treatment (*T*), whilst asymptomatic infections can be identified through screening and then receive treatment (*T*) or can recover naturally and return to the uninfected state. Depending on the vaccination strategy, individuals are vaccinated before entry into the sexually-active population, on gonorrhoea diagnosis, or on attendance for gonorrhoea screening. Upon vaccination individuals enter equivalent compartments indicated with a circumflex. Vaccine protection eventually wanes, with individuals moving into the corresponding compartments without circumflex.

### Model calibration

We calibrated the model, in a Bayesian framework, to the annual number of gonorrhoea cases among MSM in England between 2008 and 2017 recorded by the GUMCAD surveillance system.^21^ These data were considered as the observed realisations of a complex underlying unobserved Markov process including all state transitions in Figure 1. All unknown parameters were calibrated, including those governing the within-host natural history model and the risk of transmission for each sexual activity group. An analytical expression for the likelihood of the observed data given the model is not tractable, so we used a particle filter to produce an unbiased estimate.^22^ We incorporated this estimated likelihood into a particle Markov Chain Monte Carlo (pMCMC), from which we obtained a posterior sample of the model parameters given the observed data.^23,24^ Bayesian inference requires to specify prior parameter distributions, which were based on literature review where possible, or otherwise left highly uninformative to represent lack of knowledge. Full details of the modelling and a complete list of parameter values and prior distributions can be found in the appendix.

### Vaccine deployment strategies

#### We considered three realistic strategies for the vaccine deployment

“vaccination-before-entry” into the sexually active population, “vaccination-on-diagnosis” with gonorrhoea, and “vaccination-on-attendance” of a sexual health clinic for any reason, including seeking testing and treatment for symptoms or asymptomatic screening. These scenarios were compared to baseline projections in the absence of vaccination. We considered 500 hypothetical vaccine profiles ranging in protection level between 1% and 100% and lasting between 1 and 20 years. Vaccines were assumed to offer “leaky”, degree-type protection,^25^ meaning that all vaccinees were less likely, but still able, to acquire infection from contagious individuals. Vaccinated-yet-infected individuals were assumed to progress through the stages of infection in the same way as unvaccinated individuals (Figure 1). For individuals who were infected at the time of receiving the vaccine, we assumed vaccination did not alter the course of the infection, and only offered protection against subsequent acquisition of infection. Vaccine protection was assumed to wane over time, after which individuals become fully susceptible to infection once more.

We accounted for uncertainty in the natural history and infection model by basing our stochastic projections on 1,000 samples from the joint posterior distribution of the parameters. We considered varying severity of the ABR strain imported from 2020 onwards, with treatment failure rates ranging between 0% and 100%. We investigated the effect of vaccine uptake by considering scenarios of 50% to 100% acceptance among eligible MSM for each vaccine-deployment scenario.

## RESULTS

Following calibration of our epidemiological model (Figure 1), we found that the simulated epidemic curves had 99% predictive intervals that included the observed data for all years (Figure 2A). For example, our model estimated that 21,900 (95% CI 18,900-25,200) cases of gonorrhoea were reported among MSM in England in 2017, in good agreement with the 21,300 cases recorded by the GUMCAD surveillance system.^21^ In the best-case scenario, with no worsening of resistance, our model predicted 23,500 (95% CI 20,000-27,600) gonorrhoea cases in 2030. However, the global emergence and importation of an antibiotic-resistant (ABR) strain causing treatment failures would increase the projected number of cases (Figure 2B). In the worst-case scenario of systematic treatment failure for the ABR strain, our model predicted 48,500 (95% CI 23,600-80,100) gonorrhoea cases in 2030. As the frequency of treatment failure was increased, onward transmission of drug-resistant infections became more common and the predicted number of resistant cases grew.

**Figure 2.**
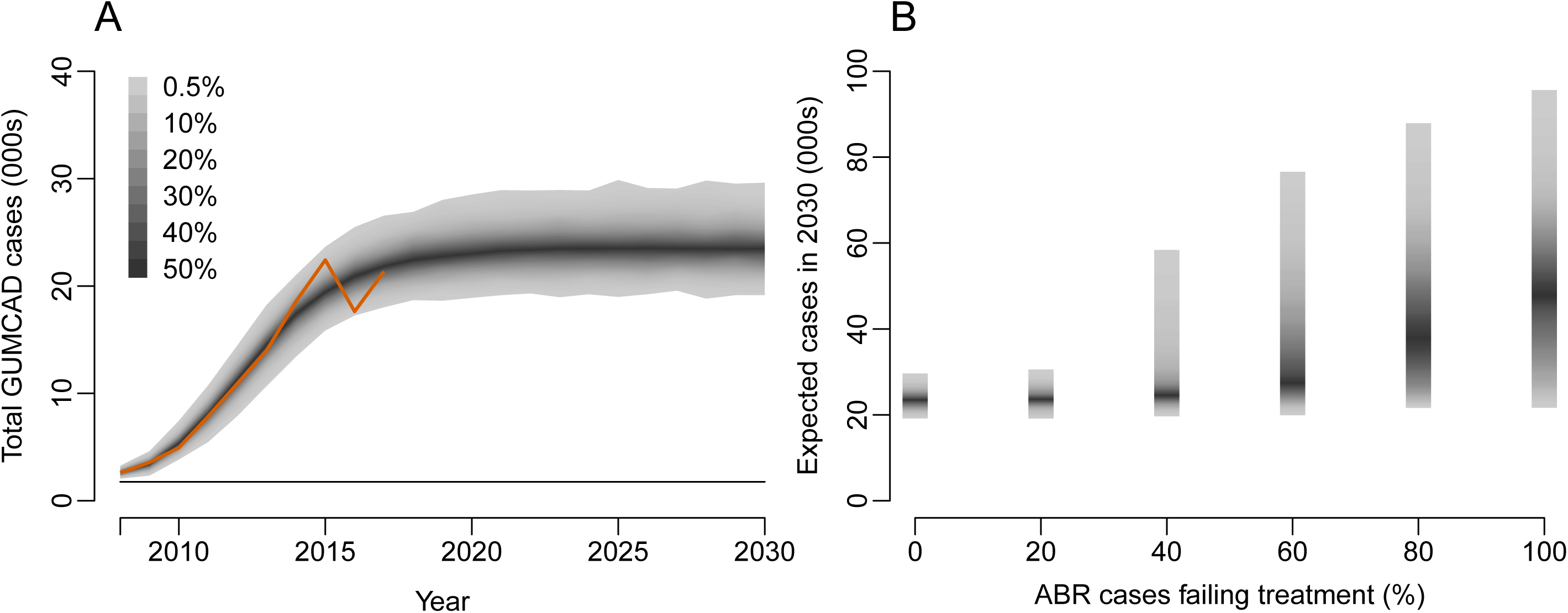
Simulated annual gonorrhoea cases between 2008 and 2030 in the absence of vaccination. The model is fitted to data in the period 2008-2017 and then projected beyond that period. **(A)** Number of cases if a novel resistant strain does not emerge. The red line depicts GUMCAD data; the horizontal line shows WHO target of 90% reduction in incidence relative to 2016. Shaded areas show 99% posterior predictive intervals, based on 1,000 simulations. **(B)** Expected number of cases in 2030 depending on the frequency of treatment failure for a novel antibiotic-resistant strain emerging in 2020.

Focussing on the worst-case scenario where treatment of ABR gonorrhoea always failed, we assessed the impact of each hypothetical vaccine profile (i.e. combination of level and duration of protection) under the three proposed deployment strategies by calculating the expected reduction in gonorrhoea cases in 2030 (Figure 3). In the vaccination-before-entry strategy, even a fully protective vaccine lasting 20 years achieved only a 34% (95% CI 17%-52%) reduction in expected cases in 2030 (Figure 3A), well below the WHO target. Vaccination-on-diagnosis with this idealised vaccine also fell short of the WHO target but achieved a much higher reduction of 92% (95% CI 82%-98%) (Figure 3B). By extending vaccination to all MSM tested for gonorrhoea (both those seeking care for symptoms and those undergoing screening in the absence of symptoms), the WHO target could be achieved using a ≥53% effective vaccine lasting ≥6 years, or equivalently a ≥70% effective vaccine lasting ≥3 years (Figure 3C). Under the vaccination-on-diagnosis and vaccination-on-attendance strategies, a vaccine lasting eight years had similar benefits to one with the same protection level lasting longer (Figure 3B,C).

**Figure 3.**
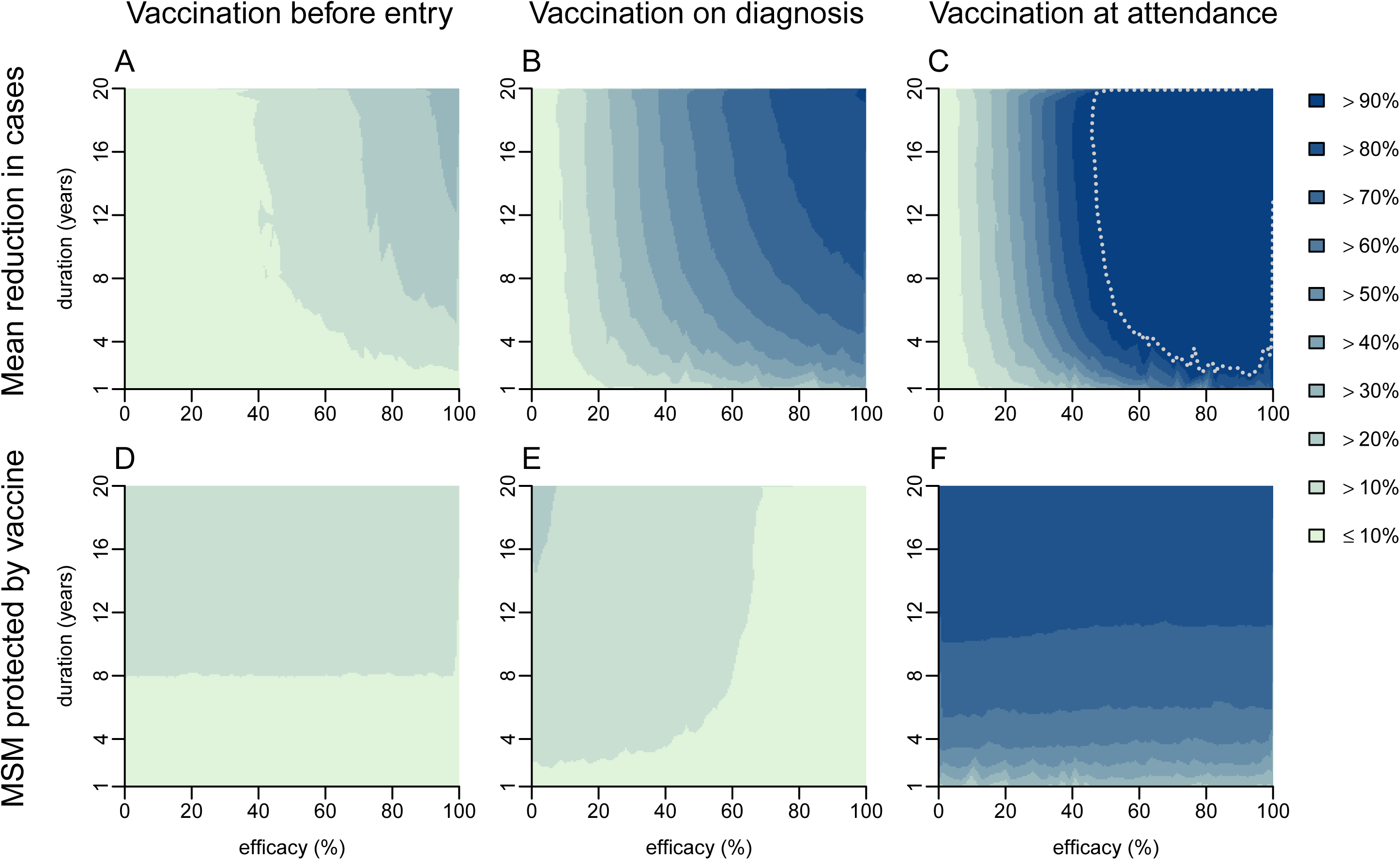
Impact of different vaccination strategies against gonorrhoea, given the emergence of a resistant strain in 2020 for which antibiotic treatment always fails. Vaccination was implemented from 2020 and administered to individuals entering the sexually active population **(A, D)**, or to patients diagnosed with gonorrhoea **(B, E)**, or to patients at clinic attendance **(C, F)**. Level (horizontal axes) and duration of protection (vertical axes) were varied. **(A, B, C)** show the mean reduction in the expected number of cases in 2030, whereas **(D, E, F)** show the proportion of MSM protected by the vaccine in 2030. Vaccine profiles (i.e. combinations of level and duration of protection) for which WHO incidence target was achieved by 2030 are highlighted with a dotted line (this is only achieved in part of **C**).

Under the vaccination-before-entry strategy, the proportion of MSM protected in 2030 was <20% irrespective of the duration of vaccine protection (Figure 3D), because only those becoming sexually active over the past 10 years had been vaccinated. The proportion of the protected population increased with the vaccine duration. Under the vaccination-on-diagnosis strategy the protected proportion was generally higher (Figure 3E) because more individuals were eligible for vaccination. Interestingly, the proportion protected decreased as vaccine-efficacy increased, because a highly protective vaccine prevented transmission which in turn reduced the number of cases treated and vaccinated alongside treatment. The vaccination-on-attendance strategy had a slightly similar pattern because it targeted everyone eligible under vaccination-on-diagnosis plus many others. The maximum proportion protected under vaccination-on-attendance was 86% (95% CI 85.8%-86.4%), if the duration of protection was 20 years (Figure 3F).

We assessed how the impact of vaccination would differ depending on the treatability of the emergent gonorrhoea ABR strain and the uptake of vaccination by eligible MSM (Figure 4). The less treatable the emergent strain of gonorrhoea, the more protective a vaccine would need to be to meet the WHO target. The vaccination-before-entry strategy was unable to achieve the WHO target, no matter how protective and long-lasting the vaccine, even in the best-case scenario of no treatment failure. The vaccination-on-diagnosis strategy met the WHO target, for a range of vaccine profiles, in scenarios where at least 30% of ABR cases remain treatable, provided almost all eligible MSM accepted the vaccine when offered and protection lasted almost 20 years (Figure 4A). With 75% vaccine uptake vaccination-on-diagnosis could not meet the target if ABR treatment failure exceeded 60% (Figure 4B). With an uptake of only 50%, the target was not met even in the best-case scenario of no treatment failure. The lower the uptake, the greater the vaccine protection level and/or duration required to achieve a given impact (Figure 4A-B). The vaccination-on-attendance strategy resulted in much higher coverage, meaning that less-protective vaccines were able to achieve the WHO target (Figure 4C). For example, in the absence of treatment failure, a 35%-protective vaccine lasting ten years could meet the WHO target, whereas, under the vaccination-on-diagnosis strategy, the vaccine would need to be 80%-protective. At the other extreme, if treatment of ABR cases always failed then the WHO target could be achieved with the vaccination-on-attendance strategy but would require for example a protection duration of six years and efficacy of 50%-75% depending on uptake (Figure 4C-E).

**Figure 4.**
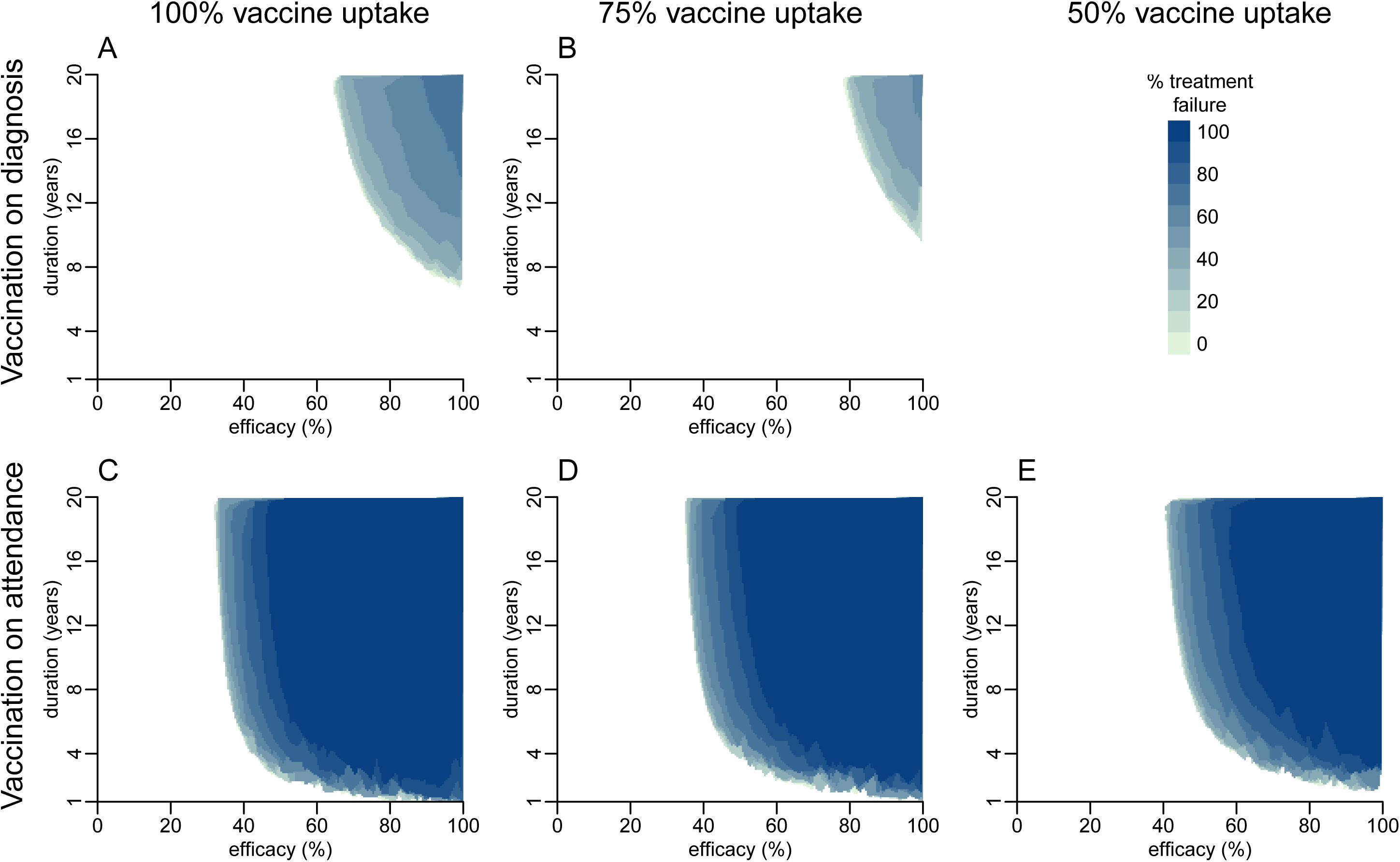
Protection level and duration of vaccine needed to reduce incidence below the WHO target by 2030 for gonorrhoea strains with varying degrees of resistance. Vaccination was implemented on diagnosis **(A, B)** or at clinic attendance **(C, D, E)**, under varying levels of vaccine uptake: 100% **(A, C)**, 75% **(B, D)** and 50% **(E)**. The vaccination-before-entry strategy is not shown as it did not achieve the WHO target for any scenario considered, and likewise for the vaccination-on-diagnosis strategy with an uptake of 50%.

Currently the only vaccine with estimated effectiveness and duration of protection against gonorrhoea is MeNZB, which offers 21%-39% protection for 2-4 years.^9^ We considered deployment of a similar vaccine using the three strategies, and compared the expected number of cases in 2030, the proportion of ABR, the proportion of protected MSM and the number of cases averted over ten years per person vaccinated (Figure 5). If there were no emergence of ABR, then a vaccine like MeNZB could have a substantial impact on the projected epidemic. Vaccination-on-diagnosis could reduce the expected incidence in 2030 by 41% (95% CI 18%-65%), from 23,500 (95% CI 20,000-27,600) to 13,900 (95% CI 8,200-18,700). Extending vaccination to all MSM tested for gonorrhoea (i.e. vaccination-on-attendance) would increase the impact to reduce incidence by 75% (95% CI 40%-98%), which was insufficient to meet the WHO target of <1,800 cases in 2030 in 85% of the simulations. The increased impact of vaccination-on-attendance came at the cost of reduced efficiency, with the mean cases averted per vaccination reduced from 0.51 (95% CI 0.32-0.75) to 0.10 (95% CI 0.07-0.13). Applying the vaccination-before-entry strategy with a MeNZB-like vaccine was only likely to achieve a 7% (95% CI 0%-23%) reduction in 2030 incidence, even without emergence of ABR.

**Figure 5.**
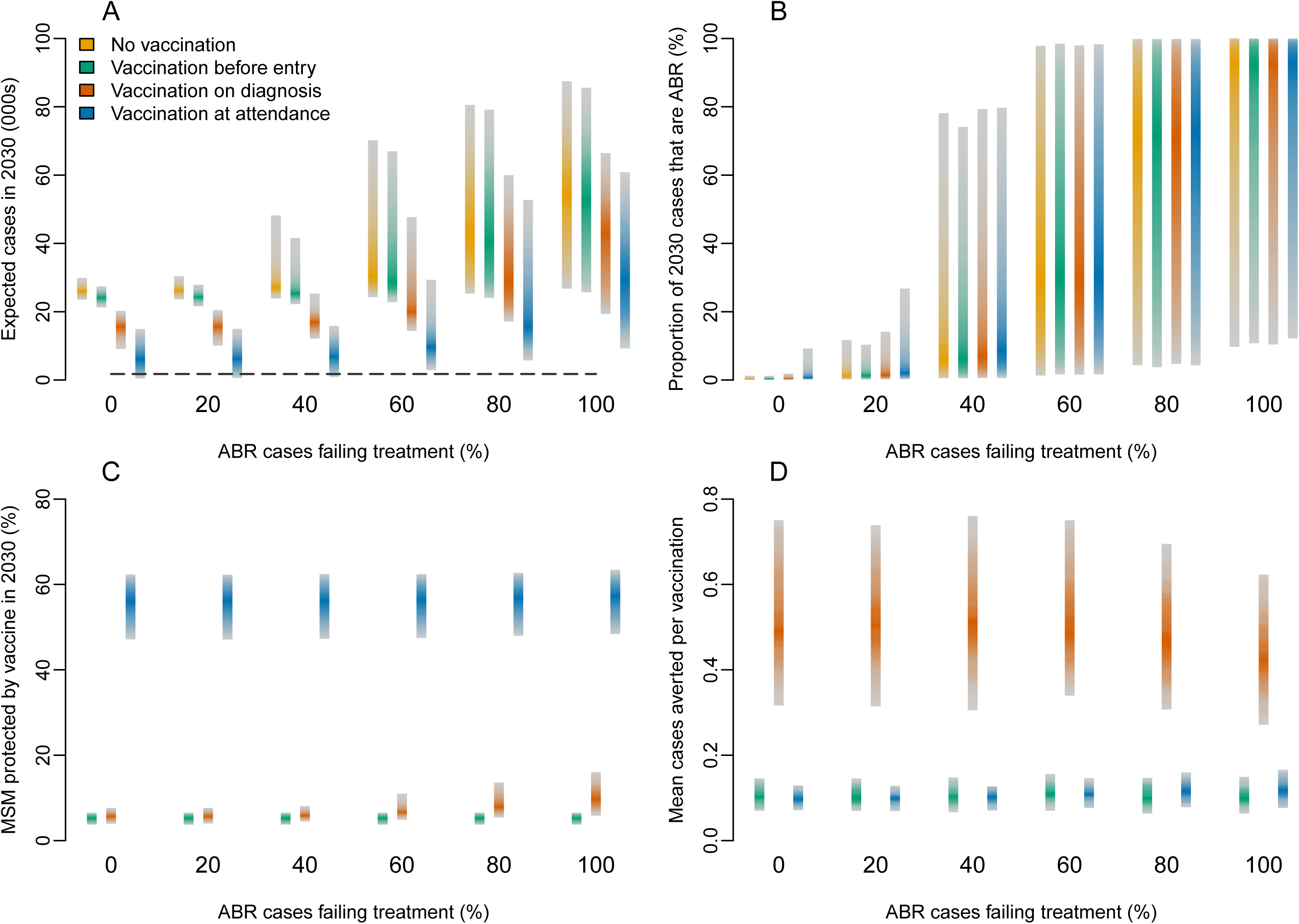
Potential impact and efficiency of the MeNZB vaccine. Vaccination was implemented before entry into the sexually active population, on diagnosis or at clinic attendance. Shaded bars depict 95% predictive interval. **(A)** Expected gonorrhoea cases in 2030, with the dashed line depicting the WHO target. **(B)** Proportion of gonorrhoea cases expected to be resistant in 2030. **(C)** Proportion of MSM protected by the vaccine in 2030. **(D)** Mean cases averted per course of vaccination, with the dashed line depicting one case averted per vaccination.

The greater the frequency at which treatment failure occurred for ABR infections, the greater the predicted incidence in 2030 and the less likely the epidemic could be controlled by a MeNZB-like vaccine (Figure 5A). Vaccination-on-attendance remained the most effective strategy, regardless of the treatability of the resistant strain. In the extreme case where ABR treatment always failed, vaccination-on-attendance with a MeNZB-like vaccine with 100% uptake reduced the expected number of cases by 45% (95% CI 18%-77%). This equated to the prevention of around 20,900 cases in 2030, reducing the expected diagnoses from 48,500 (95% CI 23,600-80,100) to 27,600 (95% CI 8,000-55,800). The proportion of cases expected to be ABR in 2030 was unaffected by vaccination, regardless of the vaccination strategy: vaccination reduced the expected diagnoses of both the resistant and susceptible strains proportionately (Figure 5B). The proportion of MSM protected by the vaccine, and the corresponding number of vaccine courses dispensed, depended more on the vaccination strategy than on the treatability of the ABR strain (Figure 5C), with vaccination-on-attendance resulting in a much higher proportion of protected MSM. Vaccination-on-diagnosis proved to be the most efficient strategy overall, with on average about one averted case for every two vaccine doses administered, which remained constant for almost all resistant strains (Figure 5D).

## DISCUSSION

We developed a stochastic transmission-dynamic model of gonorrhoea that incorporates heterogeneity in sexual behaviour and use of health services for care-seeking and screening. The model was calibrated using 10 years’ worth of data from England using Bayesian inference methods to estimate model parameters, and then used to examine the population-level impact of potential vaccines, administered via three realistic strategies. Our results show that even a partially-protective vaccine could be useful in controlling the gonorrhoea epidemic and combatting the spread of antibiotic resistance. In the absence of antibiotic treatment failure, vaccinating all MSM who attend sexual health clinics in England from 2020 onwards with a 45%-protective vaccine lasting at least four years, or a 60%-protective vaccine lasting at least two years, would be sufficient to meet the WHO target of a 90% reduction in annual incidence between 2016 and 2030.^14^ To investigate the interplay between resistance and vaccination, we considered the global emergence and importation into England of a resistant strain for which treatment sometimes fail. We made the conservative assumption that antibiotic resistant strains did not incur a fitness cost compared to antibiotic sensitive strains. In the extreme case where treatment for this new strain always fails, a 52%-protective vaccine lasting six years would be necessary to achieve the WHO target.

Imperfect vaccines are usually modelled as providing either take-type protection (where a proportion of vaccinees are fully protected and the remainder not at all), or degree-type protection (where all vaccinees have a partial reduction in the probability of infection upon exposure).^25^ It is unknown what type of protection a future gonorrhoea vaccine might provide, or indeed if it would conform to one type or another or a mixture of both, so we chose to be conservative and assume degree-type protection, which requires greater coverage to reduce the prevalence of infection than with take-type protection.^26^ Furthermore, a take-type protection vaccine is similar in result to the effect of partial uptake for a highly protective degree-type vaccine, which we considered separately (Figure 4). Uptake of vaccination is clearly a critical determinant for the success of any vaccination programme. A recent pilot programme for human papillomavirus (HPV) vaccine in UK MSM up to 45 years old recorded uptake of 45%, which is likely to be an underestimate due to incomplete recording.^27^ Therefore, the 50% uptake scenario we modelled could be considered a realistic lower bound, especially for the strategies vaccination-on-diagnosis and vaccination-on-attendance in which the vaccine would be offered to individuals likely to be concerned.

Our study is the first to consider vaccination against gonorrhoea in the context of antibiotic resistance, and to assess the potential real-world impact that could be achieved using a vaccine with protection profile similar to MeNZB. Previous modelling studies have focused on assessing the potential impact of hypothetical vaccines in small populations of either heterosexual individuals^11^ or MSM^28^. In accordance with our findings, they concluded that a vaccine of moderate level and duration of protection (60%, 10 years) could substantially reduce prevalence of infection by about 40%.^11^ Our analysis is novel in considering the effect of a vaccination on the scale of all MSM within a country, by comparing realistic deployment strategies, and by incorporating the possible global emergence of a new extensively resistant strain, as well as incorporating a statistically rigorous analysis of incidence data.

Our model assumes that sexual health services will be able to meet the additional demand caused by increasing levels of incidence and resistance. However, in recent years, access to UK sexual health services has worsened for individuals with symptoms of an acute sexually transmitted infection.^29^ This insufficient capacity in sexual health clinics could create a vicious circle, where treatment delays cause onward transmission, increased incidence, and further unmet treatment need, a situation that would be exacerbated by antibiotic resistance.^30^ A vaccine would ease this pressure by averting infections, thereby reducing demand on clinics – provided, of course, that the clinics have sufficient capacity to administer the vaccination.

Two important distinct areas for attention are the criteria on which treatment guidelines are formulated, and the need for well-designed vaccine trials with the aim of improving our understanding of the natural history of *N. gonorrhoeae*. Treatment guidelines are currently based on the proportion of diagnosed cases that are drug-resistant, with exceedance of a 5% threshold prompting changes to recommendations. However, it is important for policymakers also to consider the total number of resistant cases - especially in scenarios where resistant gonorrhoea becomes more difficult and costly to treat, as exemplified by a recent case of multi-drug resistant gonorrhoea that required three days of inpatient treatment with intravenous ertapenem.^5^

Trials of vaccine candidates need to be designed to address important gaps in knowledge, including the possibility of perverse outcomes. For example, if vaccination reduces the bacterial load then this might reduce transmission by reducing infectivity. Alternatively, it might promote transmission if it reduces the probability or severity of symptoms and thereby increases the proportion of infections that are left untreated and hence persistent.^31^ Not only could persistent infections lead to increased transmission but they may increase the probability of drug resistance evolving within asymptomatic hosts.^32^ Conversely, the resistance selection pressures could be reduced if less infections are being treated with antibiotics.^12^ The relationship between determinants of drug-resistance and antigenicity is not known and trials should monitor the diversity of lineages of *N. gonorrhoeae*. If a determinant of drug resistance is immunogenic and included in the vaccine, then this would enhance the beneficial impact of vaccination, whilst a vaccine that is more effective against drug-sensitive strains could increase the relative prevalence of resistance. Notwithstanding these interrogations and need for further research, a gonococcal vaccine could offer the hope of bringing the current epidemic of gonorrhoea under control.

