## Supplementary Material for "Vaccination against antibiotic resistant gonorrhoea for men who have sex with men in England: a modelling study"

### Supplementary Appendix

#### Materials and methods

The total number of diagnoses of gonorrhoea in MSM in England between 2008 and 2017 were extracted from the GUMCAD data published annually by PHE [1]. We adapted a previously published stochastic compartmental model of gonorrhoea to allow for sexual heterogeneity by separating the population into two groups based on sexual activity levels $g \in \{H, L\}$ [2]. The full model is illustrated in Figure S1 with notations summarised in Tables S1 and S2. An initial population of size $N$ = 600,000 was assumed with rates of population entry and exit calibrated based on ONS estimates (Figure S2). The population size was estimated based on HIV diagnoses and the European MSM Internet Survey, which suggested a UK MSM population of 3.4% [3]. The third National Survey of Sexual Attitudes and Lifestyles (Natsal-3) in which 8.4% of men reported having at least one same-sex sexual experience, with 2.6% of men having partnered with another man in the last five years [4]. This suggests a plausible range for the MSM population size of 500,000 to 1.7 million, based on a sexually active male population of 20 million in 2007 [5].

The number of MSM in each activity group is denoted $N_g(t)$ $(N_L(t) + N_H(t) = N(t))$. Individuals remain in the sexual activity group to which they are initially assigned throughout the model simulations. The proportion of the population in each sexual activity group and average number of partners of those in each activity group were calibrated according to the Natsal-3 data. The 85% of MSM having up to five partners a year were assigned to the group $L$, with the remaining 15% who had more than five partners per year assigned to group $H$.

The rate of infection for uninfected individuals depends on the probability of infection per partnership ($\beta$), the average partner change rate of the activity group ($p_g$) and the sexual mixing pattern ($\epsilon$), i.e the number of partnerships formed within and between activity groups. The parameter $\epsilon$ represents the sexual mixing coefficient and ranges from 0 to 1, representing a range of scenarios from proportionate mixing, where partners are chosen at random regardless of their sexual activity group, to assortative mixing, where partners are preferentially chosen from an individual's own sexual activity group [6].

**Table S1. Input parameter notations, descriptions and values used in simulations**

| | Parameter description | Unit | Input value(s) |
|---|---|---|---|
| $N_0$ | total population at $t_0$ | # | 600,000 |
| $q_L$ | proportion of population in group L | % | 85 |
| $\xi(t)$ | rate of new MSM entering population | year$^{-1}$ | Figure S2 |
| $\chi(t)$ | per-capita rate of MSM leaving population | year$^{-1}$ | Figure S2 |
| $p_L$ | partner-change rate in group L | year$^{-1}$ | 0.6 |
| $p_H$ | partner-change rate in group H | year$^{-1}$ | 15.6 |
| $p_E^{\text{vax}}(t)$ | proportion of new entrants that are vaccinated | % | 0-100 |
| $p_T^{\text{vax}}(t)$ | proportion of treated MSM receiving vaccine | % | 0-100 |
| $p_S^{\text{vax}}(t)$ | proportion of screened MSM receiving vaccine | % | 0-100 |
| $1/\zeta$ | expected duration of vaccine | years | 1-20 |
| $\gamma$ | vaccine-conferred reduction in susceptibility to infection | % | 1-100 |
| $\omega$ | number of imported ABR cases | year$^{-1}$ | 10 |
| $\phi$ | ABR infections for which treatment fails | % | 0-100 |

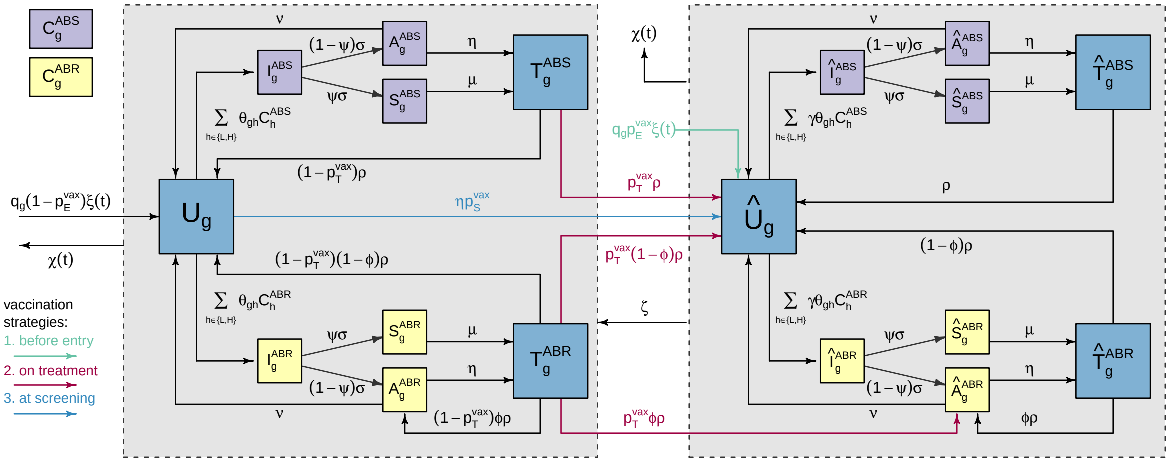

**Figure S1. Flow diagram of model compartments with rates of transition between infection states.** Uninfected individuals in group $g \in \{L, H\}$ ($U_g$) become infected with strain $s \in \{\text{ABS}, \text{ABR}\}$. Infections initially pass through an incubation period ($I_g^s$), before either developing symptoms ($S_g^s$) or remaining asymptomatic individuals ($A_g^s$). Symptomatic individuals seek care and asymptomatic individuals are identified through screening; upon which they receive treatment ($T_g^s s$). Individuals are vaccinated either before entry into the population, upon diagnosis or at sexual health clinic attendance. If the vaccine is not completely protective, vaccinated individuals can still become infected. Vaccine-conferred protection eventually wanes, which is represented by a single arrow connecting the dashed boxed rather than connecting the individual compartments.

The rate of transmission from an individual in group $h$ to an uninfected individual in group $g$ is denoted $\theta_{gh}$ and defined as follows:

$$\theta_{\mathrm{LL}}(t) = \frac{p_L\beta}{N_L(t)}\left(\epsilon + (1-\epsilon)\frac{p_L N_L(t)}{p_L N_L(t) + p_H N_H(t)}\right) \tag{S1}$$

$$\theta_{\mathrm{LH}}(t) = \frac{p_L\beta}{N_H(t)}(1-\epsilon)\frac{p_H N_H(t)}{p_L N_L(t) + p_H N_H(t)} = \frac{p_L p_H \beta(1-\epsilon)}{p_L N_L(t) + p_H N_H(t)} \tag{S2}$$

$$\theta_{\mathrm{HL}}(t) = \frac{p_H\beta}{N_L(t)}(1-\epsilon)\frac{p_L N_L(t)}{p_L N_L(t) + p_H N_H(t)} = \frac{p_L p_H \beta(1-\epsilon)}{p_L N_L(t) + p_H N_H(t)} \tag{S3}$$

$$\theta_{\mathrm{HH}}(t) = \frac{p_H\beta}{N_H(t)}\left(\epsilon + (1-\epsilon)\frac{p_H N_H(t)}{p_L N_L(t) + p_H N_H(t)}\right) \tag{S4}$$

Individuals in group $g$ are initially uninfected ($U_g$). They become infected with strain $s \in \{\mathrm{ABS}, \mathrm{ABR}\}$, denoting antibiotic susceptible (ABS) and ABR strains respectively. The model assumes that strains do not vary in transmissibility or fitness, and that the rate of infection from a contagious individual in group h to an uninfected individual in group g is $\theta_{gh}(t)$. Infected individuals initially pass through an incubation period ($I_g^s$), which they leave at rate $\sigma$. A proportion $\psi$ of those infected then go on to develop symptoms ($S_g^s$), whereas the rest remain asymptomatic ($A_g^s$). Individuals without symptoms attend sexual health screening at rate $\eta$. Gonococcal infection can occur in the rectum, pharynx and/or urethra, resulting in different rates of onward transmission and probabilities of developing symptoms [7]. We did not explicitly model separate sites of infection, therefore the rate of transmission and the likelihood of developing symptoms should be seen as an average for any infection site. Recovery from asymptomatic infection happens (either naturally or following unrelated antibiotic treatment) at rate $\nu$. The symptomatic individuals ($S_g^s$) seek care at rate $\mu$. The treated individuals recover from the infection and become uninfected again at rate $\rho$, with the exception of a proportion $\phi$ of the individuals infected with an ABR strain ($T_g^{\mathrm{ABR}}$) for whom treatment fails and who become asymptomatically infected ($A_g^{\mathrm{ABR}}$). The model assumes that no natural immunity is conferred by infection [8, 9]. The ABR strain is assumed to be imported into the highly sexually active group at rate $\omega/N_H(t)$ per person per year from time $t_{\mathrm{ABR}}$=2018 onwards.

#### Interventions

We considered three potential vaccination strategies in the model:

- Vaccination before entry into the population - where a proportion $p_E^{\text{vax}} > 0$ of adolescents are vaccinated before they become sexually active;
- Vaccination on diagnosis - where the vaccine is given to a proportion $p_T^{\text{vax}} > 0$ of MSM diagnosed with gonorrhoea;
- Vaccination at attendance - where the vaccine is given to a proportion $p_S^{\text{vax}} = p_T^{\text{vax}} > 0$ of MSM that present to sexual health services, either through screening or due to symptoms.

Vaccines were assumed to vary in efficacy, with factor $\gamma$ denoting the relative reduction in susceptibility to gonorrhoea conferred by the vaccine, which is known as degree-type protection [10]. As such, vaccinated MSM ($\hat{U}_g$) can still become infected, albeit less frequently, on sexual contact with contagious individuals ($C$). These vaccinated-yet-infected individuals progress through the stages of infection ($\hat{I}_g, \hat{S}_g, \hat{A}_g$) similarly to those who are unvaccinated. The contagious population for each strain $s$ is denoted $C_g^s = I_g^s + S_g^s + A_g^s + \hat{I}_g^s + \hat{S}_g^s + \hat{A}_g^s$. It was further assumed that under the vaccination on diagnosis strategy, treatment fails for a proportion $\phi$ of those infected with the ABR strain, who become asymptomatically infected and contagious, despite receiving the vaccine ($\hat{A}_g^{\text{ABR}}$). Vaccine protection was assumed to wane at rate $\zeta$ per year, after which time individuals become fully susceptible to infection once more.

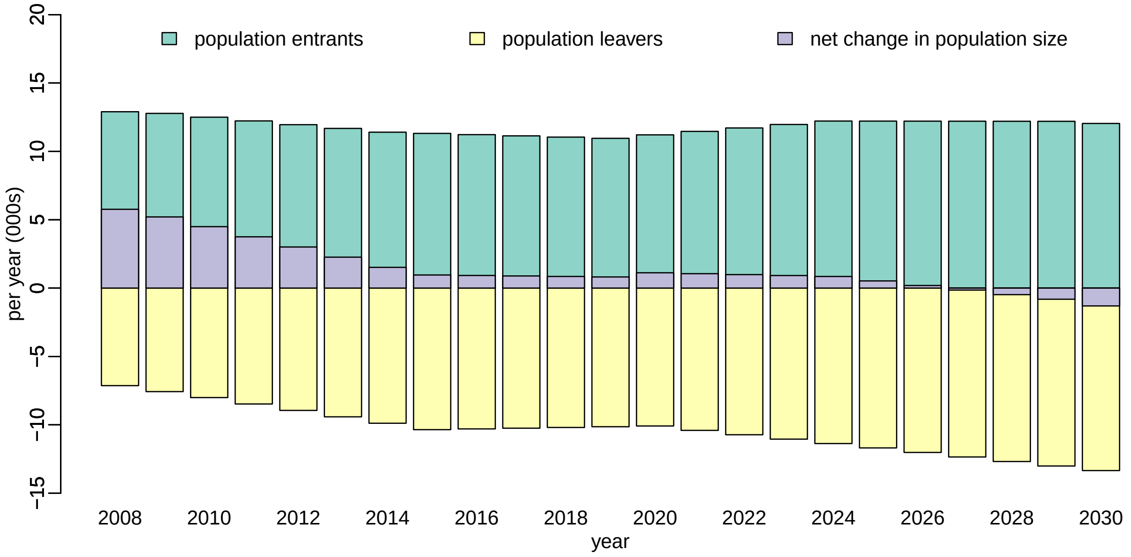

**Figure S2. Expected number of population entrants, at age 15, and leavers, at age 65, between 2008 and 2030** derived from ONS population projections assuming 3.3% of men are MSM and assuming an initial population size of 600,000

#### Model equations and stochastic simulations

A deterministic version of the model is described by the differential equations S5 - S20, in order to illustrate the model dynamics.

$$\begin{aligned}\frac{dU_H(t)}{dt} &= \big(1-p_E^{\text{vax}}(t)\big)q_H\xi(t) - \Big(\theta_{HL}C_L(t) + \theta_{HH}C_H(t) + \eta p_S^{\text{vax}}(t) + \frac{\omega(t)}{N_H(t)} + \chi(t)\Big)U_H(t) \\ &+ \nu\big(A_H^{\text{ABS}}(t) + A_H^{\text{ABR}}(t)\big) + \big(1-p_T^{\text{vax}}(t)\big)\rho\Big(T_H^{\text{ABS}}(t) + \big(1-\phi\big)T_H^{\text{ABR}}(t)\Big) + \zeta\hat{U}_H(t) \end{aligned} \tag{S5}$$

$$\begin{aligned}\frac{dU_L(t)}{dt} &= \big(1-p_E^{\text{vax}}(t)\big)q_L\xi(t) - \Big(\theta_{LL}C_L(t) + \theta_{LH}C_H(t) + \eta p_S^{\text{vax}}(t) + \chi(t)\Big)U_L(t) \\ &+ \nu\big(A_L^{\text{ABS}}(t) + A_L^{\text{ABR}}(t)\big) + \big(1-p_T^{\text{vax}}(t)\big)\rho\Big(T_L^{\text{ABS}}(t) + \big(1-\phi\big)T_L^{\text{ABR}}(t)\Big) + \zeta\hat{U}_L(t) \end{aligned} \tag{S6}$$

$$\frac{dI_H^{\text{ABR}}(t)}{dt} = \Big(\theta_{HL}C_L^{\text{ABR}}(t) + \theta_{HH}C_H^{\text{ABR}}(t) + \frac{\omega(t)}{N_H(t)}\Big)U_H(t) - \big(\sigma + \chi(t)\big)I_H^{\text{ABR}}(t) + \zeta\hat{I}_H^{\text{ABR}}(t) \tag{S7}$$

$$\frac{dI_g^s(t)}{dt} = \Big(\theta_{gL}C_L^s(t) + \theta_{gH}C_H^s(t)\Big)U_g(t) - \big(\sigma + \chi(t)\big)I_g^s(t) + \zeta\hat{I}_g^s(t); \quad \text{otherwise} \tag{S8}$$

$$\frac{dS_g^s(t)}{dt} = \psi\sigma I_g^s(t) - \big(\mu + \chi(t)\big)S_g^s(t) + \zeta\hat{S}_g^s(t); \tag{S9}$$

$$\frac{dA_g^{\text{ABS}}(t)}{dt} = \big(1-\psi\big)\sigma I_g^{\text{ABS}}(t) - \big(\nu + \eta + \chi(t)\big)A_g^{\text{ABS}}(t) + \zeta\hat{A}_g^{\text{ABS}}(t) \tag{S10}$$

$$\frac{dA_g^{\text{ABR}}(t)}{dt} = \big(1-\psi\big)\sigma I_g^{\text{ABR}}(t) - \big(\nu + \eta + \chi(t)\big)A_g^{\text{ABR}}(t) + \big(1-p_T^{\text{vax}}(t)\big)\phi\rho T_g^{\text{ABR}}(t) + \zeta\hat{A}_g^{\text{ABR}}(t) \tag{S11}$$

$$\frac{dT_g^s(t)}{dt} = \mu S_g^s(t) + \eta A_g^s(t) - \big(\rho + \chi(t)\big)T_g^s(t) + \zeta\hat{T}_g^s(t) \tag{S12}$$

$$\begin{aligned}\frac{d\hat{U}_H(t)}{dt} &= p_E^{\text{vax}}(t)q_H\xi(t) + \eta p_S^{\text{vax}}(t)U_H(t) + p_T^{\text{vax}}(t)\rho\Big(T_H^{\text{ABS}}(t) + \big(1-\phi\big)T_H^{\text{ABR}}(t)\Big) \\ &- \Big(\gamma\Big(\theta_{HL}C_L(t) + \theta_{HH}C_H(t) + \frac{\omega(t)}{N_H(t)}\Big) + \zeta + \chi(t)\Big)\hat{U}_H(t) \\ &+ \nu\Big(\hat{A}_H^{\text{ABS}}(t) + \hat{A}_H^{\text{ABR}}(t)\Big) + \rho\Big(\hat{T}_H^{\text{ABS}}(t) + \big(1-\phi\big)\hat{T}_H^{\text{ABR}}(t)\Big) \end{aligned} \tag{S13}$$

$$\begin{aligned}\frac{d\hat{U}_L(t)}{dt} &= p_E^{\text{vax}}(t)q_L\xi(t) + \eta p_S^{\text{vax}}(t)U_L(t) + p_T^{\text{vax}}(t)\rho\Big(T_L^{\text{ABS}}(t) + \big(1-\phi\big)T_L^{\text{ABR}}(t)\Big) \\ &- \Big(\gamma\big(\theta_{LL}C_L(t) + \theta_{LH}C_H(t)\big) + \zeta + \chi(t)\Big)\hat{U}_L(t) \\ &+ \nu\Big(\hat{A}_L^{\text{ABS}}(t) + \hat{A}_L^{\text{ABR}}(t)\Big) + \rho\Big(\hat{T}_L^{\text{ABS}}(t) + \big(1-\phi\big)\hat{T}_L^{\text{ABR}}(t)\Big)\end{aligned} \quad \text{(S14)}$$

$$\frac{d\hat{I}_H^{\text{ABR}}(t)}{dt} = \gamma\bigg(\theta_{HL}C_L^{\text{ABR}}(t) + \theta_{HH}C_H^{\text{ABR}}(t) + \frac{\omega(t)}{N_H(t)}\bigg)\hat{U}_H(t) - \Big(\sigma + \zeta + \chi(t)\Big)\hat{I}_H^{\text{ABR}}(t) \quad \text{(S15)}$$

$$\frac{d\hat{I}_g^s(t)}{dt} = \gamma\Big(\theta_{gL}C_L^s(t) + \theta_{gH}C_H^s(t)\Big)\hat{U}_g(t) - \Big(\sigma + \zeta + \chi(t)\Big)\hat{I}_g^s(t); \quad \text{otherwise} \quad \text{(S16)}$$

$$\frac{d\hat{S}_g^s(t)}{dt} = \psi\sigma\hat{I}_g^s(t) - \big(\mu + \zeta + \chi(t)\big)\hat{S}_g^s(t) \quad \text{(S17)}$$

$$\frac{d\hat{A}_g^{\text{ABS}}(t)}{dt} = \big(1-\psi\big)\sigma\hat{I}_g^{\text{ABS}}(t) - \big(\nu + \eta + \zeta + \chi(t)\big)\hat{A}_g^{\text{ABS}}(t) \quad \text{(S18)}$$

$$\frac{d\hat{A}_g^{\text{ABR}}(t)}{dt} = p_T^{\text{vax}}\phi\rho T_g^{\text{ABR}}(t) + \big(1-\psi\big)\sigma\hat{I}_g^{\text{ABR}}(t) - \big(\nu + \eta + \zeta + \chi(t)\big)\hat{A}_g^{\text{ABR}}(t) + \phi\rho\hat{T}_g^{\text{ABR}}(t) \quad \text{(S19)}$$

$$\frac{d\hat{T}_g^s(t)}{dt} = \mu\hat{S}_g^s(t) + \eta\hat{A}_g^s(t) - \big(\rho + \zeta + \chi(t)\big)\hat{T}_g^s(t) \quad \text{(S20)}$$

We used a stochastic version of the model described in Equations S5 - S20, time-discretised using the Gillespie tau-leap method with steps of 1 day. The process was initialised on 31 December 2007, with $A_g^{\mathrm{ABS}}(0)$ asymptomatic infections in each sexual activity group $g \in \{L, H\}$ and the remainder of the population uninfected. Simulation proceeded through iterations of the steps below for each day. Since the model is stochastic, the outbreak can become extinct even when its basic reproduction number is $>1$. Transition variables $b_1^g, b_2^g, d_1^g, \ldots, d_{32}^g$ are drawn from the following distributions:

$$b_1^g \sim \mathrm{Poisson}\Big(q_g\big(1-p_E^{\mathrm{vax}}\big)\xi(t)\Big) \tag{S21}$$

$$b_2^g \sim \mathrm{Poisson}\Big(q_g p_E^{\mathrm{vax}}\xi(t)\Big) \tag{S22}$$

$$d_1^H, \ldots, d_4^H \sim \mathrm{Multinom}\Bigg(U_H(t),\ \theta_{HL}C_L^{\mathrm{ABS}}(t) + \theta_{HH}C_H^{\mathrm{ABS}}(t),\ \theta_{HL}C_L^{\mathrm{ABR}}(t) + \theta_{HH}C_H^{\mathrm{ABR}}(t) + \frac{\omega(t)}{N_H(t)},\ \eta p_S^{\mathrm{vax}}(t),\ \chi(t)\Bigg) \tag{S23}$$

$$d_1^L, \ldots, d_4^L \sim \mathrm{Multinom}\Bigg(U_L(t),\ \theta_{LL}C_L^{\mathrm{ABS}}(t) + \theta_{LH}C_H^{\mathrm{ABS}}(t),\ \theta_{LL}C_L^{\mathrm{ABR}}(t) + \theta_{LH}C_H^{\mathrm{ABR}}(t),\ \eta p_S^{\mathrm{vax}}(t),\ \chi(t)\Bigg) \tag{S24}$$

$$d_5^g, d_6^g, d_7^g \sim \mathrm{Multinom}\Big(I_g^{\mathrm{ABS}}(t),\ \psi\sigma,\ \big(1-\psi\big)\sigma,\ \chi(t)\Big) \tag{S25}$$

$$d_8^g, d_9^g, d_{10}^g \sim \mathrm{Multinom}\Big(I_g^{\mathrm{ABR}}(t),\ \psi\sigma,\ \big(1-\psi\big)\sigma,\ \chi(t)\Big) \tag{S26}$$

$$d_{11}^g, d_{12}^g \sim \mathrm{Multinom}\big(S_g^{\mathrm{ABS}}(t),\ \mu,\ \chi(t)\big) \tag{S27}$$

$$d_{13}^g, d_{14}^g \sim \mathrm{Multinom}\big(S_g^{\mathrm{ABR}}(t),\ \mu,\ \chi(t)\big) \tag{S28}$$

$$d_{15}^g, d_{16}^g, d_{17}^g \sim \mathrm{Multinom}\big(A_g^{\mathrm{ABS}}(t),\ \nu,\ \eta,\ \chi(t)\big) \tag{S29}$$

$$d_{18}^g, d_{19}^g, d_{20}^g \sim \mathrm{Multinom}\big(A_g^{\mathrm{ABR}}(t),\ \nu,\ \eta,\ \chi(t)\big) \tag{S30}$$

$$d_{21}^g, d_{22}^g, d_{23}^g \sim \mathrm{Multinom}\Big(T_g^{\mathrm{ABS}}(t),\ \big(1-p_T^{\mathrm{vax}}\big)\rho,\ p_T^{\mathrm{vax}}\rho,\ \chi(t)\Big) \tag{S31}$$

$$d_{24}^g, \ldots, d_{28}^g \sim \mathrm{Multinom}\Big(T_g^{\mathrm{ABR}}(t),\ \big(1-p_T^{\mathrm{vax}}\big)\big(1-\phi\big)\rho,\ \big(1-p_T^{\mathrm{vax}}\big)\phi\rho,\ p_T^{\mathrm{vax}}\big(1-\phi\big)\rho,\ p_T^{\mathrm{vax}}\phi\rho,\ \chi(t)\Big) \tag{S32}$$

$$d_{29}^H, \ldots, d_{32}^H \sim \mathrm{Multinom}\Bigg(\hat{U}_H(t),\ \gamma\Big(\theta_{HL}C_L^{\mathrm{ABS}}(t) + \theta_{HH}C_H^{\mathrm{ABS}}(t)\Big),\ \gamma\Big(\theta_{HL}C_L^{\mathrm{ABR}}(t) + \theta_{HH}C_H^{\mathrm{ABR}}(t) + \frac{\omega(t)}{N_H(t)}\Big),\ \zeta,\ \chi(t)\Bigg) \tag{S33}$$

$$d_{29}^L, \ldots, d_{32}^L \sim \mathrm{Multinom}\Bigg(\hat{U}_L(t),\ \gamma\Big(\theta_{LL}C_L^{\mathrm{ABS}}(t) + \theta_{LH}C_H^{\mathrm{ABS}}(t)\Big),\ \gamma\Big(\theta_{LL}C_L^{\mathrm{ABR}}(t) + \theta_{LH}C_H^{\mathrm{ABR}}(t)\Big),\ \zeta,\ \chi(t)\Bigg) \tag{S34}$$

$$d_{33}^g, \ldots d_{36}^g \sim \mathrm{Multinom}\Big(\hat{I}_g^{\mathrm{ABS}}(t),\ \psi\sigma,\ \big(1-\psi\big)\sigma,\ \zeta,\ \chi(t)\Big) \tag{S35}$$

$$d_{37}^g, \ldots, d_{40}^g \sim \mathrm{Multinom}\Big(\hat{I}_g^{\mathrm{ABR}}(t),\ \psi\sigma,\ \big(1-\psi\big)\sigma,\ \zeta,\ \chi(t)\Big) \tag{S36}$$

$$d_{41}^g, d_{42}^g, d_{43}^g \sim \mathrm{Multinom}\big(\hat{S}_g^{\mathrm{ABS}}(t),\ \mu,\ \zeta,\ \chi(t)\big) \tag{S37}$$

$$d_{44}^g, d_{45}^g, d_{46}^g \sim \mathrm{Multinom}\big(\hat{S}_g^{\mathrm{ABR}}(t),\ \mu,\ \zeta,\ \chi(t)\big) \tag{S38}$$

$$d_{47}^g, \ldots, d_{50}^g \sim \mathrm{Multinom}\big(\hat{A}_g^{\mathrm{ABS}}(t),\ \nu,\ \eta,\ \zeta,\ \chi(t)\big) \tag{S39}$$

$$d_{51}^g, \ldots, d_{54}^g \sim \mathrm{Multinom}\big(\hat{A}_g^{\mathrm{ABR}}(t),\ \nu,\ \eta,\ \zeta,\ \chi(t)\big) \tag{S40}$$

$$d_{55}^g, d_{56}^g, d_{57}^g \sim \mathrm{Multinom}\Big(\hat{T}_g^{\mathrm{ABS}}(t),\ \rho,\ \zeta,\ \chi(t)\Big) \tag{S41}$$

$$d_{58}^g, \ldots, d_{61}^g \sim \mathrm{Multinom}\Big(\hat{T}_g^{\mathrm{ABR}}(t),\ \big(1-\phi\big)\rho,\ \phi\rho,\ \zeta,\ \chi(t)\Big) \tag{S42}$$

For ease of notation, we have expressed the equations above using the rates we have previously defined, rather than their respective probabilities. Therefore, for example, $d \sim \mathrm{Multinom}(\mathrm{S}, \underline{\mathrm{r}})$ should be understood as $d \sim \mathrm{Multinom}(\mathrm{S}, 1 - \mathrm{e}^{-\underline{\mathrm{r}}})$.

The compartments of the model were updated as follows:

$$U_g(t+1) := U_g(t) + b_1^g - d_1^g - d_2^g - d_3^g - d_4^g + d_{15}^g + d_{18}^g + d_{21}^g + d_{24}^g + d_{31}^g \quad \text{(S43)}$$

$$I_g^{\mathrm{ABS}}(t+1) := I_g^{\mathrm{ABS}}(t) + d_1^g - d_5^g - d_6^g - d_7^g + d_{35}^g \quad \text{(S44)}$$

$$I_g^{\mathrm{ABR}}(t+1) := I_g^{\mathrm{ABR}}(t) + d_2^g - d_8^g - d_9^g - d_{10}^g + d_{39}^g \quad \text{(S45)}$$

$$S_g^{\mathrm{ABS}}(t+1) := S_g^{\mathrm{ABS}}(t) + d_5^g - d_{11}^g - d_{12}^g + d_{42}^g \quad \text{(S46)}$$

$$S_g^{\mathrm{ABR}}(t+1) := S_g^{\mathrm{ABR}}(t) + d_8^g - d_{13}^g - d_{14}^g + d_{45}^g \quad \text{(S47)}$$

$$A_g^{\mathrm{ABS}}(t+1) := A_g^{\mathrm{ABS}}(t) + d_6^g - d_{15}^g - d_{16}^g - d_{17}^g + d_{49}^g \quad \text{(S48)}$$

$$A_g^{\mathrm{ABR}}(t+1) := A_g^{\mathrm{ABR}}(t) + d_9^g - d_{18}^g - d_{19}^g - d_{20}^g + d_{25}^g + d_{53}^g \quad \text{(S49)}$$

$$T_g^{\mathrm{ABS}}(t+1) := T_g^{\mathrm{ABS}}(t) + d_{11}^g + d_{16}^g - d_{21}^g - d_{22}^g - d_{23}^g + d_{56}^g \quad \text{(S50)}$$

$$T_g^{\mathrm{ABR}}(t+1) := T_g^{\mathrm{ABR}}(t) + d_{13}^g + d_{19}^g - d_{24}^g - d_{25}^g - d_{26}^g - d_{27}^g - d_{28}^g + d_{60}^g \quad \text{(S51)}$$

$$\hat{U}_g(t+1) := \hat{U}_g(t) + b_2^g + d_3^g + d_{22}^g + d_{26}^g - d_{29}^g - d_{30}^g - d_{31}^g - d_{32}^g + d_{47}^g + d_{51}^g + d_{55}^g + d_{58}^g \quad \text{(S52)}$$

$$\hat{I}_g^{\mathrm{ABS}}(t+1) := \hat{I}_g^{\mathrm{ABS}}(t) + d_{29}^g - d_{33}^g - d_{34}^g - d_{35}^g - d_{36}^g \quad \text{(S53)}$$

$$\hat{I}_g^{\mathrm{ABR}}(t+1) := \hat{I}_g^{\mathrm{ABR}}(t) + d_{30}^g - d_{37}^g - d_{38}^g - d_{39}^g - d_{40}^g \quad \text{(S54)}$$

$$\hat{S}_g^{\mathrm{ABS}}(t+1) := \hat{S}_g^{\mathrm{ABS}}(t) + d_{33}^g - d_{41}^g - d_{42}^g - d_{43}^g \quad \text{(S55)}$$

$$\hat{S}_g^{\mathrm{ABR}}(t+1) := \hat{S}_g^{\mathrm{ABR}}(t) + d_{37}^g - d_{44}^g - d_{45}^g - d_{46}^g \quad \text{(S56)}$$

$$\hat{A}_g^{\mathrm{ABS}}(t+1) := \hat{A}_g^{\mathrm{ABS}}(t) + d_{34}^g - d_{47}^g - d_{48}^g - d_{49}^g - d_{50}^g \quad \text{(S57)}$$

$$\hat{A}_g^{\mathrm{ABR}}(t+1) := \hat{A}_g^{\mathrm{ABR}}(t) + d_{27}^g + d_{38}^g - d_{51}^g - d_{52}^g - d_{53}^g - d_{54}^g + d_{59}^g \quad \text{(S58)}$$

$$\hat{T}_g^{\mathrm{ABS}}(t+1) := \hat{T}_g^{\mathrm{ABS}}(t) + d_{41}^g + d_{48}^g - d_{55}^g - d_{56}^g - d_{57}^g \quad \text{(S59)}$$

$$\hat{T}_g^{\mathrm{ABR}}(t+1) := \hat{T}_g^{\mathrm{ABR}}(t) + d_{44}^g + d_{52}^g - d_{58}^g - d_{59}^g - d_{60}^g - d_{61}^g \quad \text{(S60)}$$

#### Bayesian Inference

We considered the GUMCAD recorded cases ($Y_D(t)$, t = 2008, ..., 2017) as the observed realisations of an underlying unobserved Markov process: the total number of diagnosed gonococcal infections ($Z_D(t)$) (Figure S3). We assumed that 90% of all diagnoses are recorded by GUMCAD, with a 10% margin of under-reporting - consistent with evidence of the proportion of gonorrhoea diagnoses made in a General Practice setting, and therefore not reported to GUMCAD [11].

$$Y_D(t) \sim \mathcal{N}(0.9 Z_D(t), 0.05 Z_D(t)) \quad \text{(S61)}$$

Additionally, to ensure that the rate at which MSM attend sexual health screening reflects the most recent data, we considered the number of sexual health clinic attendances in 2017 ($Y_A(t)$, t = 2017) as the observed realisation of an underlying unobserved variable - the total number of MSM attending sexual health services ($Z_A(2017)$), again allowing for a 10% margin of under-reporting.

$$Y_A(2017) \sim \mathrm{Poisson}(0.9 Z_A(2017)) \quad \text{(S62)}$$

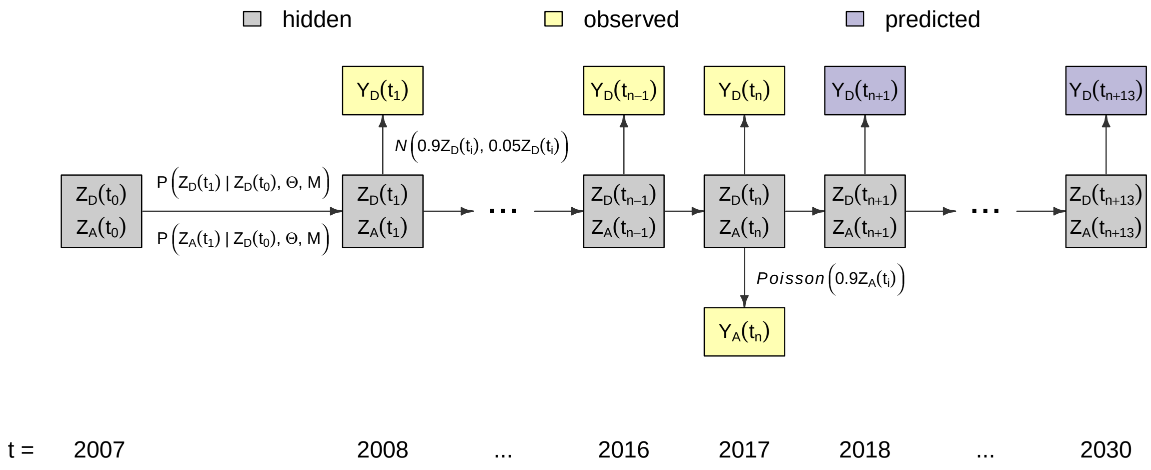

**Figure S3. Illustrative diagram of the partially observed Markov process** The total incidence of gonorrhoea infections, $Z_D(t)$, and annual sexual health clinic attendances, $Z_A(t)$ are unobserved Markovian time series, updated via the model of infection, M, with parameters $\Theta$. The number of GUMCAD recorded cases, $Y_D(t)$, and recorded attendances, $Y_A(t)$ are the observed realisations of the underlying hidden processes.

An expression for the likelihood of the observed data given the model is not analytically tractable, so we used a particle filter to compute an unbiased estimate. This estimated likelihood was then incorporated into a Monte Carlo Markov Chain (MCMC) algorithm, which was used to sample from the joint posterior distribution of the parameters given the observed data [12].

As the number of parameters increases, so does the difficulty of exploring the joint posterior space. To be parsimonious we reduced the number of model parameters by making the initial number of symptomatic cases, $S_L(2007)$ and $S_H(2007)$, to be zero.. After a few simulated days these variables quickly reach the stochastic values implied by the model and parameters. Therefore, the first few days of each simulation can be thought of as the equivalent to a Markov Chain convergence or burn-in step.

We implemented the model fitting using a pMCMC algorithm from the R package pomp [13], which we modified to enable parallel computation. The estimation of the likelihood was based on 1,000 particles, which we found to be sufficiently robust to produce consistent estimates of the likelihood. Four pMCMC with dispersed starting points were run for 110,000 iterations, with the first 10% discarded as burn-in. The chains were compared using the R package coda [14]; the multivariate Gelman-Rubin diagnostic was $\leq$1.1 for all inferred parameters, suggesting that they appear to have converged on the same posterior distribution [15]. For robustness, we combined the posterior samples from all four chains and ensured all parameters had an effective sample size greater than 200.

#### Prior distributions of parameters

Performing Bayesian inference requires the specification of plausible parameter priors. For parameters whose values are uncertain ($\epsilon$, $\frac{A_L^{\text{ABS}}(0)}{N_L}$, $\frac{A_H^{\text{ABS}}(0)}{N_H}$, $\beta, \psi$) we adopted highly uninformative uniform priors (Table S2). The remaining four parameters $\nu, \sigma, \mu$ and $\rho$ were assigned informative priors based on literature review, as summarised in Table S2.

The length of time for which asymptomatic gonorrhoea infection persists has not been well measured. Duration of carriage likely depends on the site of infection, with estimates varying widely from 12 weeks at the pharynx, up to a year for rectal infection [16, 17]. In a study of 18 asymptomatic men infected with (urethral) gonorrhoea, no infections were cleared in the 165 days from diagnosis to treatment [18]. Modelling studies commonly asume the average duration of untreated infection is 6 months [6, 19, 20]. This is supported by evidence from genomic data, where bacterial genomes isolated from known contact pairs had a maximum time to most recent common ancestor of 8 months [21]. We therefore assigned the duration of carriage, $\frac{1}{\nu}$, a prior corresponding to a mean duration of carriage between three months and one year with 95% prior weight.

The UK national guidelines on safer sex advice recommend MSM test for gonorrhoea annually, and that highly sexually active MSM get tested every three months [22, 23]. However, in a social media survey of 2,668 MSM in Scotland, Wales and Ireland, two-thirds reported STI testing less than once a year [24]. We assigned a uniform distribution to the rate of screening, $\eta$ with 95% prior weight between 0.4 and 4. This corresponds to a range of scenarios: from testing at quarterly intervals, in adherence with the BASHH guidelines, through to the situation described by [24]. The lower bound for $\eta$ was calculated by setting the probability of testing less than once a year equal to $\frac{2}{3}$ under an exponential distribution with rate $\eta$, so that $\mathcal{P}[T > 1] = e^{-\eta} = \frac{2}{3}$, and solving to find $\eta = -\ln\frac{2}{3} = 0.4$.

The length of time spent in the incubation, symptomatic, and treatment stages of infection are observed to be short [25–27]. We set priors accordingly: the mean duration of incubation, $\frac{1}{\sigma}$, was assumed to be between three and eight days; the mean duration of symptomatic infection (before seeking care), $\frac{1}{\mu}$, was assumed to be between a day and two weeks; and the mean duration of treatment, $\frac{1}{\rho}$, was assumed to be between five and nine days, in accordance with the BASHH recommendation to abstain from sex for seven days after treatment [28].

**Table S2. Calibrated parameter notations, descriptions, prior and posterior distributions**

| | Parameter description | Unit | Prior distribution | Prior mean (95% CI) | Posterior mean (95% CrI) |
|---|---|---|---|---|---|
| $\epsilon$ | level of assortativity in sexual mixing | - | $\mathcal{U}[0,1]$ | 0.5 (0.025, 0.975) | 0.43 (0.06, 0.74) |
| $\frac{A_L^{\mathrm{ABS}}(0)}{N_L}$ | asymptomatic prevalence in group L at $t_0$ | % | $\mathcal{U}[0,10]$ | 5 (0.25, 9.75) | 0.9 (0.2, 1.7) |
| $\frac{A_H^{\mathrm{ABS}}(0)}{N_H}$ | asymptomatic prevalence in group H at $t_0$ | % | $\mathcal{U}[0,10]$ | 5 (0.25, 9.75) | 1.0 (0.4, 1.7) |
| $\beta$ | probability of transmission per partnership | % | $\mathcal{U}[0,100]$ | 50 (2.5, 97.5) | 58.5 (28.3, 96.5) |
| $\psi$ | infections that become symptomatic | % | $\mathcal{U}[0,100]$ | 50 (2.5, 97.5) | 24.5 (13.3, 35.2) |
| $1/\sigma$ | average duration incubation period | days | $\mathrm{InvGamma}(17,4.6)$ | 4.7 (3.0, 8.0) | 4.6 (3.0, 7.6) |
| $1/\nu$ | average duration of asymptomatic carriage | days | $\mathrm{InvGamma}(8,0.29)$ | 157 (87, 364) | 174 (95, 443) |
| $\eta$ | rate of screening when asymptomatic | year$^{-1}$ | $\mathrm{U}[0.4,4]$ | 2.2 (0.49, 3.91) | 0.45 (0.44, 0.46) |
| $1/\mu$ | average time to symptomatics seeking care | days | $\mathrm{InvGamma}(3,45.6)$ | 2.7 (1.1, 12.9) | 2.8 (1.2, 14.7) |
| $1/\rho$ | average time to recovery following treatment | days | $\mathrm{InvGamma}(101,0.52)$ | 7.0 (5.8, 8.5) | 6.9 (5.8, 8.6) |

#### Estimation of model parameters

The posterior estimate of the probability of transmission per partnership per year $\beta$ was 59% with a wide range of credibly values from 28% to 97%. The parameter was positively correlated with the level of assortativity in sexual mixing $\epsilon$, which was estimated to be between 0.06 and 0.75 with 95% credibility. This corresponds to the trade-off required to maintain a force of infection compatible with the observed data, as described by equations S1-4.

The model predicts that 25% (95% CrI: 13%, 35%) of infections become symptomatic, and that this proportion ($\psi$) is strongly negatively correlated with the rate of asymptomatic screening ($\eta$). This corresponds to the tradeoff between duration of carriage, $\frac{1}{\nu+\eta}$, and the proportion of infections entering the carriage state $(1-\psi)$

The parameters corresponding to the time spent in the incubation, symptomatic, and treatment stages of infection ($\sigma, \mu, \rho$) had posterior distributions that closely matched their prior distributions, suggesting that the priors were appropriate and that there is little additional information on these parameters to be gleaned from this data set. These parameters did not show a strong correlation with any others; as expected given the short duration of the incubation, symptomatic and treatment stages variation in their parameters makes little difference to the dynamics of infection.

The posterior distribution of $\nu$ has a slightly lower mean than the prior, implying a longer mean duration of carriage: 174 (95% CrI: 95, 443) days compared to 157 (95% CrI: 87, 364). The prior and posterior credible intervals intersect to a large extent so there is not significant evidence of a departure from the prior based on the data.

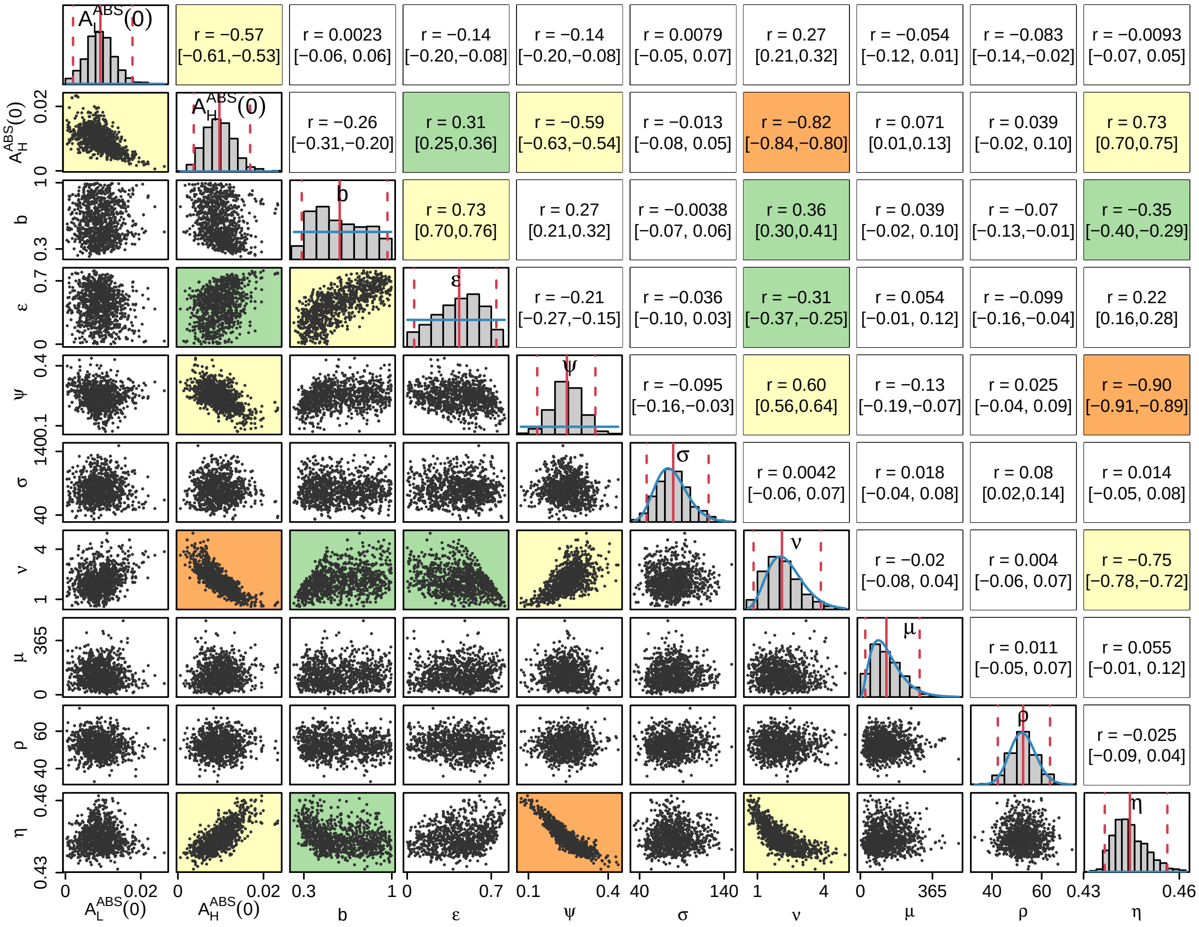

**Figure S4. Posterior distributions of parameters.** The diagonal plots show histograms of the posterior distributions for all sampled parameters. The blue lines show prior distributions, and the red lines indicate posterior mean and 95% credible intervals. The plots below the diagonal show scatter plots based on 1,000 samples from the posterior, illustrating the relationships between pairs of estimated parameters. An orange background indicates a correlation higher than 0.8, a yellow background indicates a correlation between 0.5 and 0.8, a green background indicates a correlation between 0.3 and 0.5, and a white background indicates a correlation less than 0.3. The plots above the diagonal show the corresponding correlation coefficients with the 95% credible intervals in parentheses.

#### Overview of simulation analysis

In order to assess the necessary quality of a gonococcal vaccine, and how it would be best deployed, we considered 1,000 vaccine profiles (i.e. combinations of level and duration of protection), selected using Latin Hypercube Sampling, which ranged in efficacy from 1-100% and offered protection lasting between 1 and 20 years. For each vaccine profile we performed 100 Monte Carlo simulations of the epidemic between 2008 and 2030. we repeated the analysis for each of the ABR strains of varying controllability, under each of the three vaccination strategies.

We assessed the potential impact of a MeNZB-like vaccine by considering 100 vaccine profiles ranging in efficacy from 21%-39% conferring protection from infection for two to four years. We repeated the analysis for varying levels of vaccine coverage to assess how the level of uptake achieved impacts the success of each strategy.
